# Human endogenous retrovirus envelope proteins alter extracellular vesicle cellular interactions and biodistribution

**DOI:** 10.64898/2026.04.30.722014

**Authors:** Zach Troyer, Miranda Soumakis, Erin N. Shirk, Olesia Gololobova, Sarah Marquez, Martina Fabiano, Bianca Pachane, Taekyung Ryu, Chan-Hyun Na, Natalie Castell, Isabella Baumann, Suzanne Queen, Joseph L. Mankowski, Kenneth W. Witwer

## Abstract

Extracellular vesicles (EVs) are versatile therapeutic candidates due to biological roles in intercellular communication and amenability to bioengineering. Compared with lipid nanoparticles (LNPs), native or surface-modified EVs may have favorable immunogenicity and biodistribution profiles. However, when administered intravenously (IV), EVs are rapidly cleared and accumulate mostly in the liver and spleen. With the goal of modifying EV biodistribution, we engineered EVs to display the human endogenous retrovirus (HERV) envelope glycoprotein Syncytin-1, an SLC1A5-binding fusogenic viral protein essential for syncytiotrophoblast formation in pregnancy. Here, we comprehensively characterize engineered Syncytin-1+ EVs, examine their interactions with cells *in vitro*, and assay biodistribution, immunogenicity, and pharmacokinetics *ex vivo* and *in vivo* in non-human primates. IV-administered Syncytin-1+ EVs are well tolerated, persist in the blood stream, and have altered organ biodistribution compared with unmodified EVs, suggesting therapeutic potential of Syncytin-1+ EVs at specific sites.

## Introduction

Extracellular vesicles (EVs) are lipid bilayer-delimited nanoparticles released constitutively by cells from the plasma membrane (ectosomes) and multivesicular bodies (exosomes).^1–3^ EVs remove excess cellular contents, initiate signaling cascades, deliver antigens to immune cells, transfer transmembrane proteins, and induce pro- and anti-inflammatory responses.^2^ Multiple clinical trials of EVs are currently underway, investigating their utility as diagnostics and therapeutics.^4^ Although most EV studies have focused on presumed intercellular transfer of luminal content, EV-cell fusion now appears to be a relatively rare event, and specific loading of luminal content is challenging.^5,6^ EV-cell fusion and other intercellular signaling strategies involving EVs are likely mediated by surface components, which can be relatively easily engineered through pre- and post-production strategies.^7–12^ EV surface engineering may also enhance biodistribution/selective retention advantages of EVs over lipid nanoparticles (LNPs), taking advantage of protein-protein interactions.^13–15^

Numerous surface engineering strategies have been used to improve EV-cell fusion and achieve selective retention of EVs by specific cell types. For fusion, vesicular stomatitis virus G protein (VSV-G) or another viral fusion protein is expressed on the surface of EVs^16–19^. VSV-G is highly fusogenic and promiscuous, binding the widely expressed low-density lipoprotein receptor (LDL-R)^20^. However, the high immunogenicity of viral proteins is a barrier to clinical use^19,21^. For selective retention, often referred to as targeting or homing, EVs have been engineered to display ligands for selected cell/tissue-specific receptors. This includes EV modification with Sialyl Lewis-X (sLeX) carbohydrates^9^, modified antibody-derivatives targeting the mannose receptor^22^ (CD206) and CD19^23^, PD-1^23,24^, tetraspanins^25^, IL-3^26^, RVG^27^ and RGD^28^ peptides, and nucleic acid aptamers^29^.

For either selective retention or fusion to occur in the mammalian organism, the EV must first reach its cellular target.^25,30^ However, studies of both rodents and non-human primates (NHPs) report that the majority of IV-administered EVs are rapidly filtered out of blood by liver and spleen, in addition to clearance by phagocytic immune cells.^21,31–36^ Furthermore, although EVs are generally considered to have low immunogenicity, repeated administration of exogenous EVs to NHPs in resulted in antigenicity and accelerated blood clearance (ABC).^36^ Strategies to slow clearance or avoid immunogenicity include masking the EV with “don’t-eat-me signals” or abundant blood proteins such as albumin and using patient-derived EVs.^21,37,38^

In this study, we engineer EVs to carry human endogenous retrovirus (HERV) envelope proteins Syncytin-1^39^, Syncytin-2^40^, or HERV-K 108 Envelope^41^ to enhance cell binding and fusion and assess their interactions with cells *in vitro*. We also examine the pharmacokinetics, biodistribution, and immunogenicity of Syncytin-1+ EVs *in vivo* in NHPs. We reasoned that, since the coding sequences for these proteins have been endogenized during primate evolution and the proteins are expressed at different stages of development, they may be less immunogenic than exogenous viral proteins, recognized as “self.”^42^ We further reasoned that these proteins might alter the expected biodistribution of EVs by binding specific receptors that are less widely expressed than LDL-R. As an example, Syncytin-1 is found in hominoids and some other primates, and its fusogenicity is essential for syncytiotrophoblast formation in pregnancy.^43,44^ Syncytin-1 recognizes SLC1A5 on recipient cells.^45^ In addition to trophoblasts, SLC1A5 is expressed on various other cells, including barrier endothelial cells and with strong expression in kidney and bladder.^46,47^ Our results suggest that, in comparison with unmodified EVs, Syncytin-1+ EVs have increased circulatory half-life, reduced immune cell interactions, reduced liver accumulation, increased kidney accumulation, and reduced immunogenicity. These results support further development of HERV envelope-bearing EVs to target specific organ diseases.

## Results

### Selection and expression of functional HERV envelope proteins

For EV engineering, Syncytin-1 (HERV-W Envelope), Syncytin-2 (HERV-FRD Envelope) and HERV-K 108 Envelope were selected since each 1) has an intact open reading frame in humans, 2) produces a functional protein, and 3) has at least one known cellular receptor. HEK293T cells transfected with plasmids encoding each of the three candidates formed syncytia (Figure 1A), although syncytium formation was less pronounced in the presence of HERV-K 108. Western blots confirmed expression of the three HERV proteins (Figure 1B). Most of the detected HERV-K 108 Envelope in the cell lysate was immature, with the surface (SU) and transmembrane (TM) subunits remaining connected, with a smaller amount of mature protein (solitary TM subunit) resulting from furin processing.

**Figure 1.**
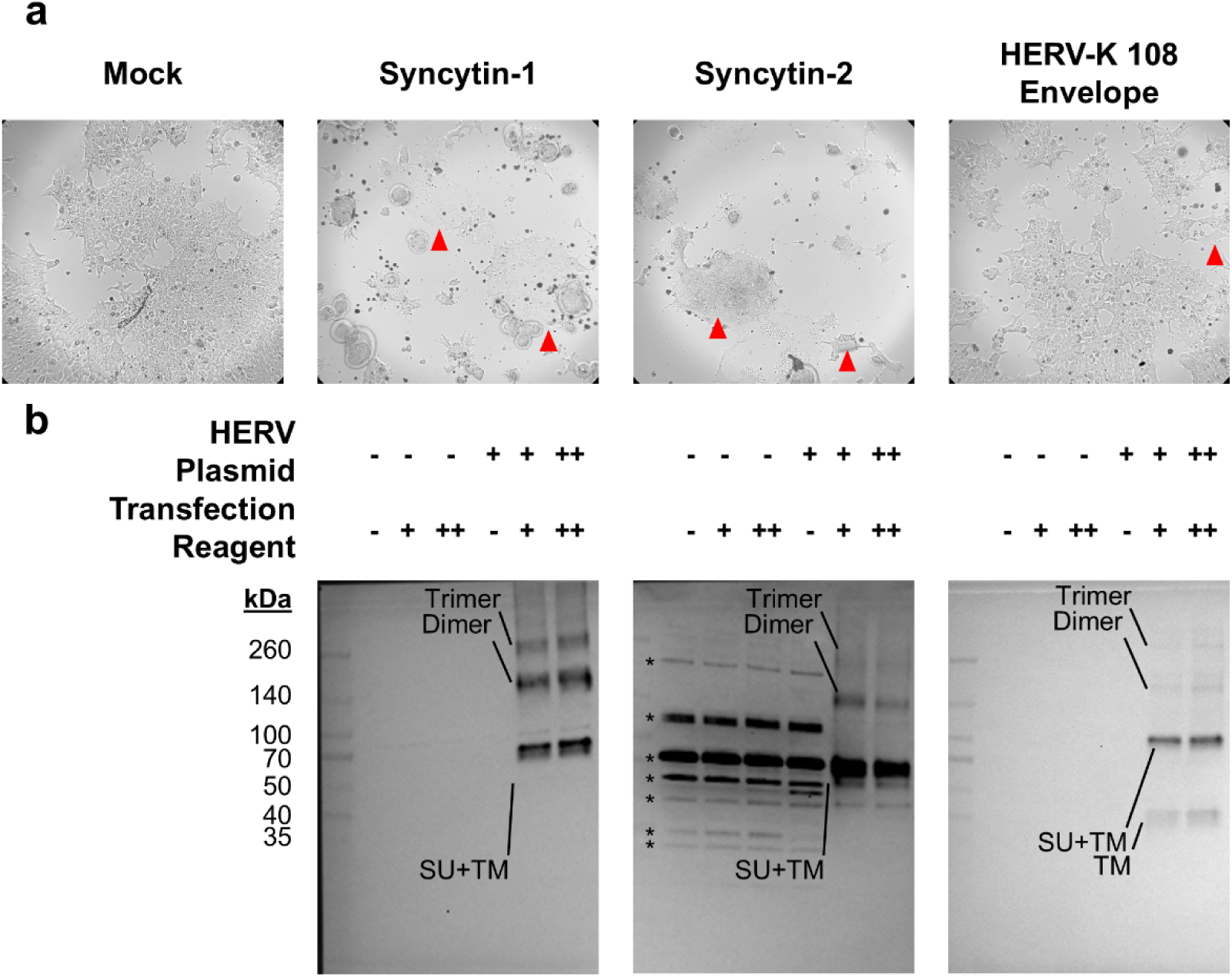
Cellular overexpression of human endogenous retrovirus (HERV) envelope proteins. (a) Brightfield microscope images of HERV envelope transfected HEK293T cells. Red triangles mark cellular syncytia. (b) Immunoblot analysis of HERV envelope transfected HEK293T lysates. 3.5 µg of lysate was loaded per well. For each HERV envelope, bands corresponding to the full-length protein, envelope dimers or trimers, or furin cleavage products are indicated by labels. Asterisks denote non-specific bands.

### Extracellular vesicle engineering and production

EVs were produced from plasmid-transfected Expi293F cells, a suspension-adapted HEK293T derivative. Similar to HEK293Ts, envelope-expressing Expi293F cells showed morphology changes such as clumping and membrane ruffling (Figure 2A). EVs were separated from 72-hour conditioned media using ultrafiltration and size-exclusion chromatography (SEC). By nanoflow cytometry, engineered and native EVs had a similar size profile, although particle number concentrations of engineered EVs were lower (Figure 2B). Western blots confirmed several EV markers and depletion of calnexin, a cellular marker (Figure 2C). Syncytin-1, Syncytin-2, and HERV-K 108 Envelope were detected in engineered EVs. Transmission electron microscopy (TEM) revealed the presence of EVs in all conditions with typical cup-shaped morphology expected of negative-stained TEM (Figure 2D). EVs or other membrane structures with irregular morphology were also present in conditions of HERV envelope overexpression. To further validate the incorporation of HERV envelope proteins on the EV surface, immunogold TEM was performed using gold-conjugated anti-HERV envelope antibodies (Figure 2E). Immunogold TEM micrographs revealed EVs with surface-clustered gold particles when the antibody matched the overexpression condition. Together, these data support successful display of mature HERV envelope proteins on engineered EVs.

**Figure 2.**
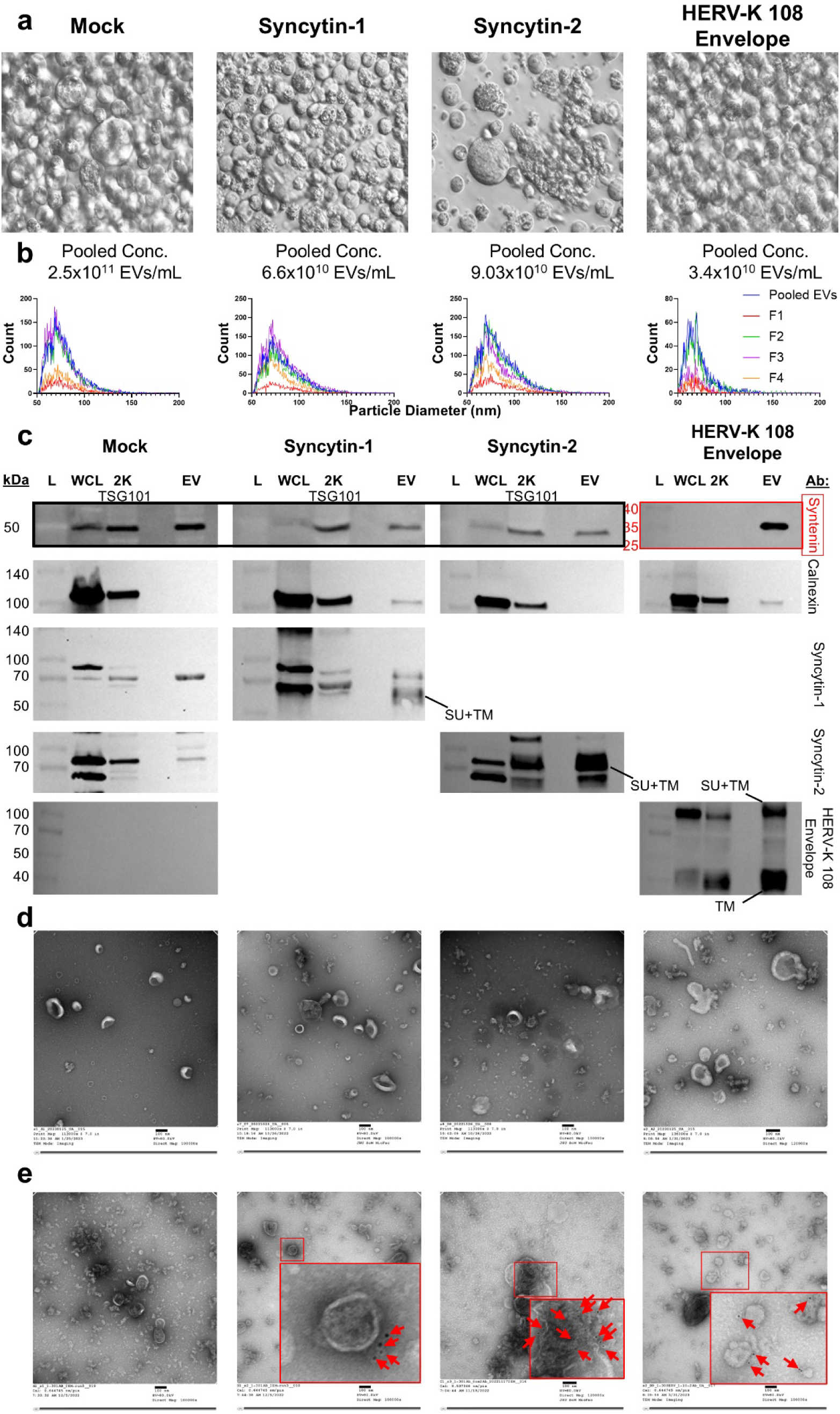
Production and characterization of HERV envelope bearing extracellular vesicles (EVs). (a) Brightfield microscope images of HERV envelope transfected Expi293F cells. (b) Nanoflow cytometry plots showing the size distribution of HERV envelope EVs derived from Expi293F cells. (c) Immunoblot analysis of protein ladder (L), whole cell lysate (WCL), 2000g (2K) pellet, and pooled EVs from derived from HERV envelope transfected Expi293F cells. TSG101 was used as an EV marker for the mock, Syncytin-1, and Syncytin-2 conditions; Syntenin-1 was used for the HERV-K 108 Envelope condition. For HERV envelope proteins, bands corresponding to the full-length protein or furin cleavage products (if detectable by the antibody used) are indicated by labels. (d) Transmission electron microscopy (TEM) micrographs of HERV envelope EVs. The scale bar represents 100 nm. (e) Immunogold TEM micrographs of HERV envelope EVs using antibodies targeting HERV envelope proteins. The inset magnified boxes identify EVs for which 6 nm gold particle-labeled anti-HERV envelope secondary antibodies clustered at the EV surface, identified by red arrows. The scale bar represents 100 nm.

### Cell and EV distribution of mature and immature envelope protein

As mentioned previously, HERV envelope proteins include a surface (SU) domain (containing a receptor-binding domain (RBD)) and a transmembrane (TM) domain that anchors the protein and effects fusion after binding.^48^ The furin protease cleaves the immature (whole, SU+TM) protein into SU and TM (mature protein). To understand the distribution of immature and mature protein in cells and EVs, we used immunoblotting with antibodies that detect only immature protein (Syncytin-1 and -2) or immature and mature HERV-K 108. For HERV-K108, EVs, in contrast with cell lysate, contained more mature protein (TM subunit) than immature protein, consistent with natural envelope maturation as the protein leaves the cells on the EV surface. To also detect mature Syncytin-1 and -2, we HA-tagged the C-terminus of the protein. Cells transfected with the HA-tagged envelope constructs had similar morphology to those transfected with the non-HA tagged versions, and EVs from these cells had similar size distributions (Supplemental Figure S1A, B). Syncytin-1+ EVs contained less immature protein than the cell lysate, and immature Syncytin-1 was also found in non-EV (protein-enriched) SEC fractions (Figure 1C, Supplemental Figure S1C). Syncytin-2 and HERV-K 108 EVs contained more immature protein than the cell lysate, suggesting preferential export, and Syncytin-2 was also found in protein SEC fractions. The majority of Syncytin-1 on EVs was in the mature form, while cells primarily carried the immature form (Supplemental Figure S1C). For Syncytin-2, EVs had roughly equal amounts of mature and immature form, while immature protein predominated in cells.

### HERV envelopes alter EV interactions with cells *in vitro*

To allow EV tracking, we next engineered cells to contain the PalmGRET dual fluorescence/luminescence (EGFP and NanoLuc luciferase) reporter^49^ along with HERV envelopes or VSV-G (Figure 3A). By nanoflow cytometry, around 30% - 50% of produced EVs were positive for PalmGRET (Figure 3B). EGFP+ EVs across all conditions tended to be slightly larger than EGFP-EVs (∼89.3 nm vs. 72 nm diameter), whether because of preferential packaging into larger EVs or incorporation increasing EV size. TEM revealed no major morphological differences with incorporation of PalmGRET (Figure 3C); VSV-G protein was visible coating the EV surface in this condition. Western blot also confirmed co-enrichment of EGFP from PalmGRET with HERV envelope proteins and the EV marker Syntenin-1 in pooled EVs (Figure 3D).

**Figure 3.**
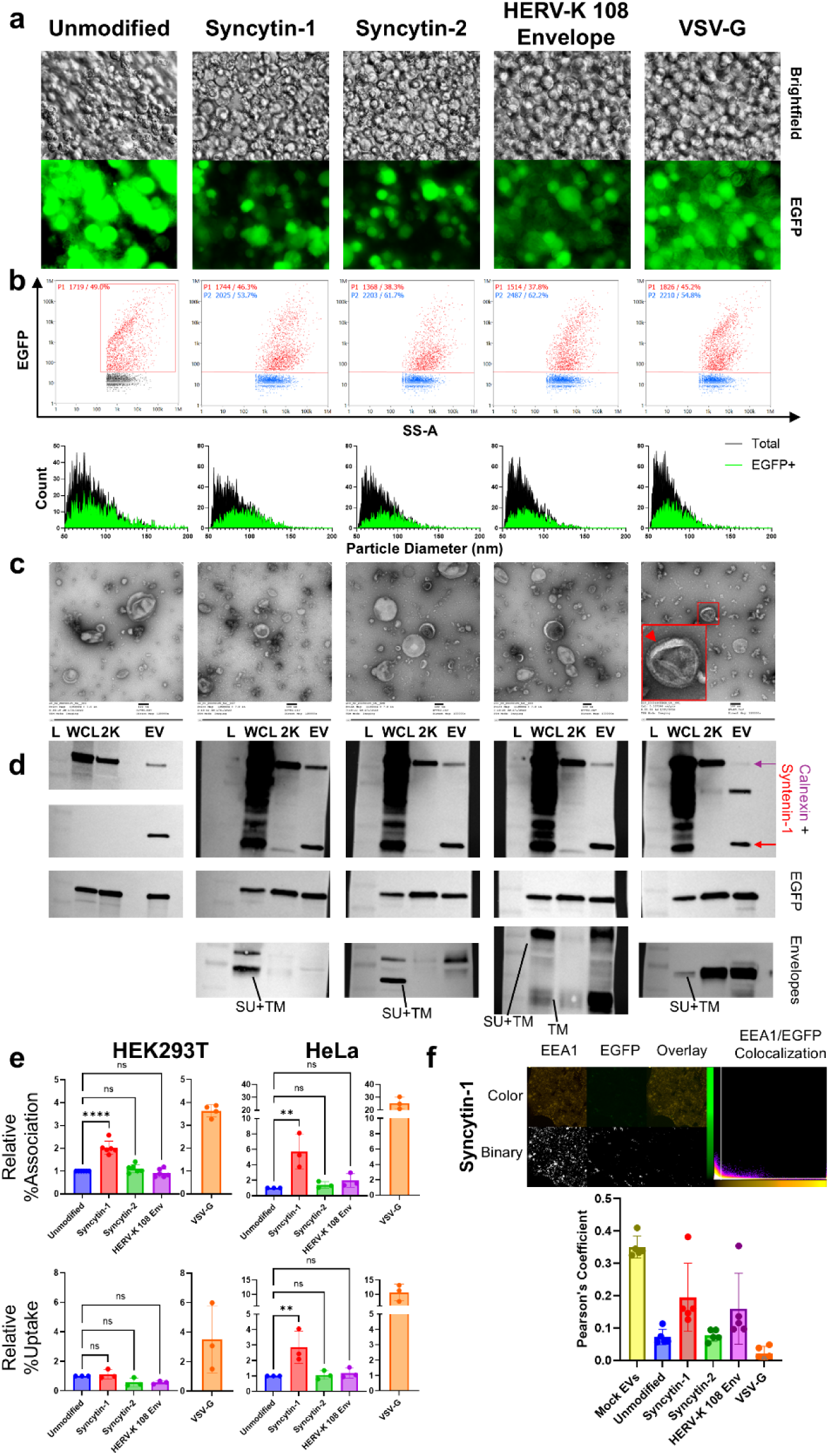
Characterization of PalmGRET/HERV envelope EVs and in vitro interactions. (a) Brightfield (top panels) and fluorescence (bottom panels) microscopy images of Expi293F cells transfected with envelope proteins (HERV and VSV-G) and the EV tracking molecule PalmGRET. (b) Nanoflow cytometry plots showing the %EGFP+ EV population (top panels) and the size distribution (lower panels) of the total and EGFP+ EV populations. (c) Transmission electron microscopy (TEM) micrographs of various PalmGRET EVs. The scale bar represents 100 nm. (d) Immunoblot analysis of ladder (L), whole cell lysate (WCL), 2000g (2K) pellet, and pooled EVs from PalmGRET/HERV envelope transfected Expi293F cells. For HERV envelope proteins, bands corresponding to the full-length protein or furin cleavage products are labeled. (e) Bar graphs representing fold change in EV association (top panels) or EV uptake (bottom panels) of PalmGRET/HERV envelope EVs with HEK293T (left panels) or HeLa (right panels) cells. VSV-G EVs are separately represented as a positive control. Bars represent mean fold-change %association or %uptake values, normalized to the unmodified EV control, from 6 (%association/HEK293T) or 3 independent biological replicates. Error bars represent standard deviation. Statistical testing was performed by one-way ANOVA with Dunnett’s correction for multiple comparisons; differences with p<0.05 were considered to be statistically significant. ns, P > 0.05; *, P ≤ 0.05; **, P ≤ 0.01; ***, P ≤ 0.001; ****, P ≤ 0.0001. (f) Representative confocal microscopy images (63X magnification) of HEK293T cells after incubation with Syncytin-1/PalmGRET EVs. Image was adjusted by increasing brightness by 40% and lowering contrast by 40%. Cells were stained with antibodies or fluorescent moieties to identify endosomes (EEA1-PE), nuclei (Hoechst 33342 – not visible), and cytoskeletal actin (Phalloidin-647 – not visible). Colocalization of green (EGFP+ EVs) and orange (EEA1+ endosomes) pixels was assessed to determine EV colocalization/escape from endosomes and quantified using Pearson correlation. Lower panel: Bar graph quantifying the correlation between the EV and endosomal fluorescent signals, for different conditions. Bars represent the mean Pearson’s coefficient of 5 confocal images; error bars represent standard deviation. Higher Pearson’s coefficient mean higher correlation and less EV endosomal escape; lower coefficient values mean less correlation and more efficient EV endosomal escape.

To assess EV-cell interactions, PalmGRET EVs with or without HERV envelopes were added to HEK293T and HeLa cells for 6 hours. Using NanoLuc luciferase assays and VSV-G+/PalmGRET EVs as a control, we measured association (i.e., all EV-cell interactions including internalization and cell surface binding) and uptake (i.e., internalized EVs that are resistant to trypsin digestion). Syncytin-1+ EVs had a statistically significant, 2-fold higher association with HEK293T cells than unmodified EVs (Figure 3E, p<0.0001). However, cellular uptake was not significantly different, suggesting major differences in surface interactions. Association and uptake of PalmGRET EVs with HEK293Ts was also observed by confocal microscopy (Figure 3F, Supplemental Figure S2). As a rough measure of endosomal escape, EV signal (green) was assessed for colocalization with the endosomal marker EEA1 (orange). All EVs showed some level of endosomal escape in HEK293Ts, with the VSV-G positive control EVs having the most (Supplemental Figure S2). In HeLa cells, Syncytin-1+ EVs had approximately 6-fold higher cellular association compared with unmodified EVs (Figure 3E, p=0.0038). Unlike in HEK293Ts, Syncytin-1+ EVs also had 3-fold higher uptake than unmodified EVs (Figure 3E, p=0.01). In both cell lines, Syncytin-2+ and HERV-K 108 Envelope+ EVs did not have significantly different association or uptake levels than unmodified EVs at 6h post-EV addition. VSV-G EVs had the highest levels of association and uptake with HEK293T and HeLa cells compared with unmodified and HERV envelope modified EVs.

### HERV envelopes reduce EV-cell interactions in NHP blood *ex vivo*

Hypothesizing that HERV envelope decoration might affect EV association with cells in blood, we introduced 1×10^9^ HERV-modified PalmGRET EVs into pigtailed macaque blood for 30 minutes *ex vivo* (Figure 4A). EV spike-in did not influence the frequency of PBMC subtypes (Supplemental Figure S3A) as determined by flow cytometry (Supplemental Figure S3B). Cell association trends were similar for all modified EVs and consistent with our previous findings^36^: CD20+ B cells had the highest levels of EV association, followed by monocytes (CD14^Hi^ and CD14^Lo^) and CD4+ T cells; CD8+ T cells and CD159a+ NK cells did not have detectable EV interactions (Figure 4B). However, all EVs with viral envelope proteins (both HERVs and VSV-G) had less cell association than unmodified EVs (Figure 4B, Total PBMC panels). This was the case across PBMC types, but greatest for CD14^Hi^ monocytes, followed by CD14^Lo^ monocytes and CD4+ T cells. EV-B cell associations were also diminished, but not as strongly. A similar interaction trend to that detected by flow cytometry was also observed in luciferase-based assay with PBMCs, further validating the results (Supplemental Figure S3C).

**Figure 4.**
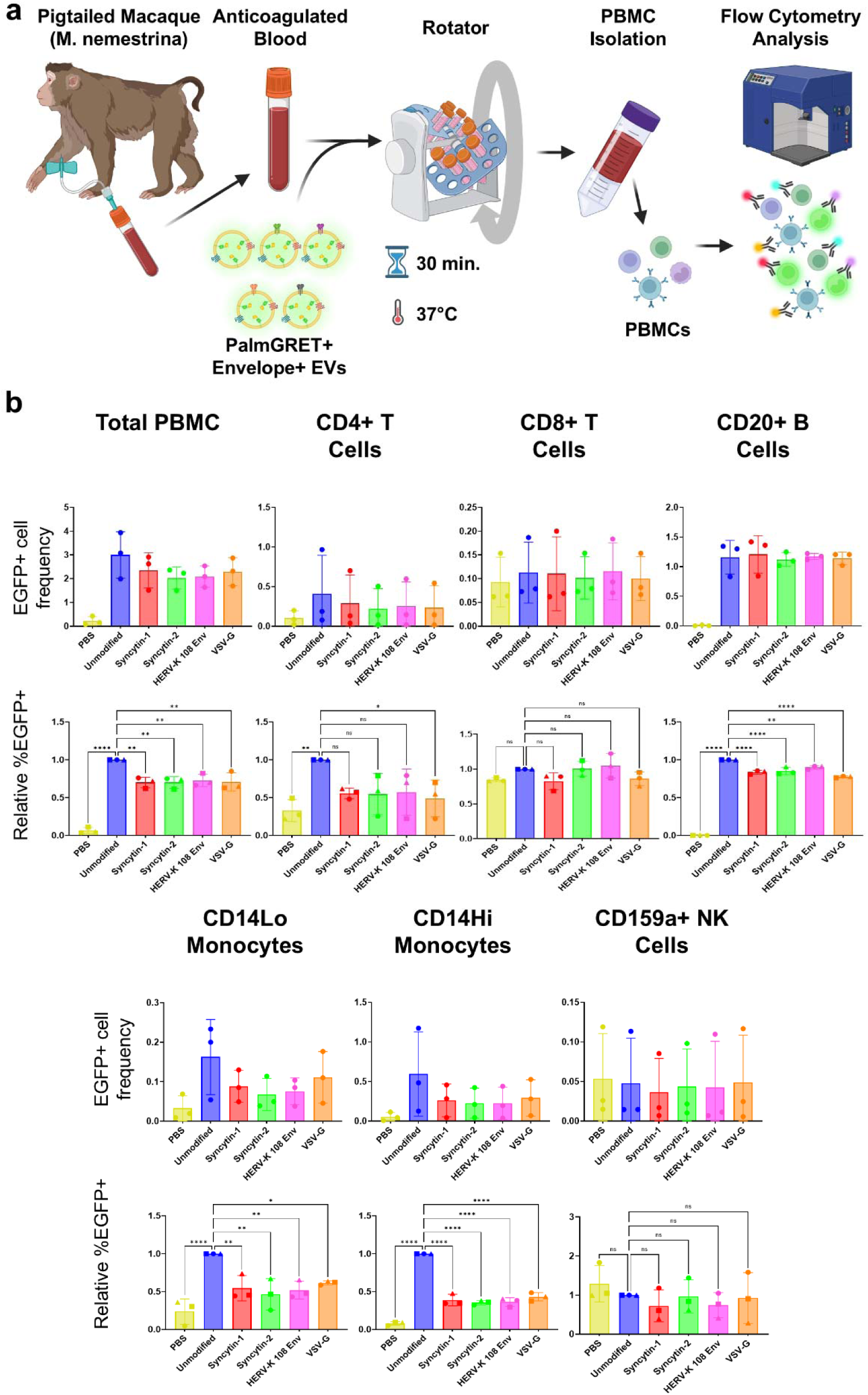
PalmGRET/HERV envelope EV interactions with non-human primate (NHP) blood ex vivo. (a) Schematic representation of the experimental workflow to collect NHP blood, mix it with modified PalmGRET EVs, and measure EV interactions with peripheral blood mononuclear cells (PBMCs) using flow cytometry. (b) Bar graphs representing the interaction of PalmGRET/HERV envelope EVs with various PBMC subsets. Bars either represent mean quantity of EGFP+ (EV-interacting) cells (top panels) or mean fold change in %EGFP+ cells (bottom panels) normalized to the unmodified PalmGRET EV condition, from 3 independent biological replicates using blood from different NHP donors. Error bars represent standard deviation. Statistical testing was performed by one-way ANOVA with Dunnett’s correction for multiple comparisons; differences with p<0.05 were considered to be statistically significant. ns, P > 0.05; *, P ≤ 0.05; **, P ≤ 0.01; ***, P ≤ 0.001; ****, P ≤ 0.0001.

### Blood cell association, pharmacokinetics, and CSF distribution of intravenously injected Syncytin-1+ EVs

Based on the *in vitro* and *ex vivo* findings described above, we chose Syncytin-1+ EVs for *in vivo* NHP biodistribution studies. Four doses of Syncytin-1+ EVs, separated by two weeks, were administered intravenously to two adult female pigtailed macaques (Figure 5A). The first three doses (4×10^10^ EVs) were Syncytin-1+ EVs only, while the final dose (1.1×10^11^ EVs) was Syncytin-1+/PalmGRET EVs to allow tracking in collected biofluids and solid tissues. Neither subject showed signs of physical or behavioral stress after EV administrations. By flow cytometry (Figure 5B) and complete blood counts (Supplemental Figure S4A), blood cell makeup did not change substantially over the course of treatment. One subject had an apparent increase in neutrophils and a corresponding decrease in CD4+ and CD8+ T cells after the first two EV doses, but not for the last two doses.

**Figure 5.**
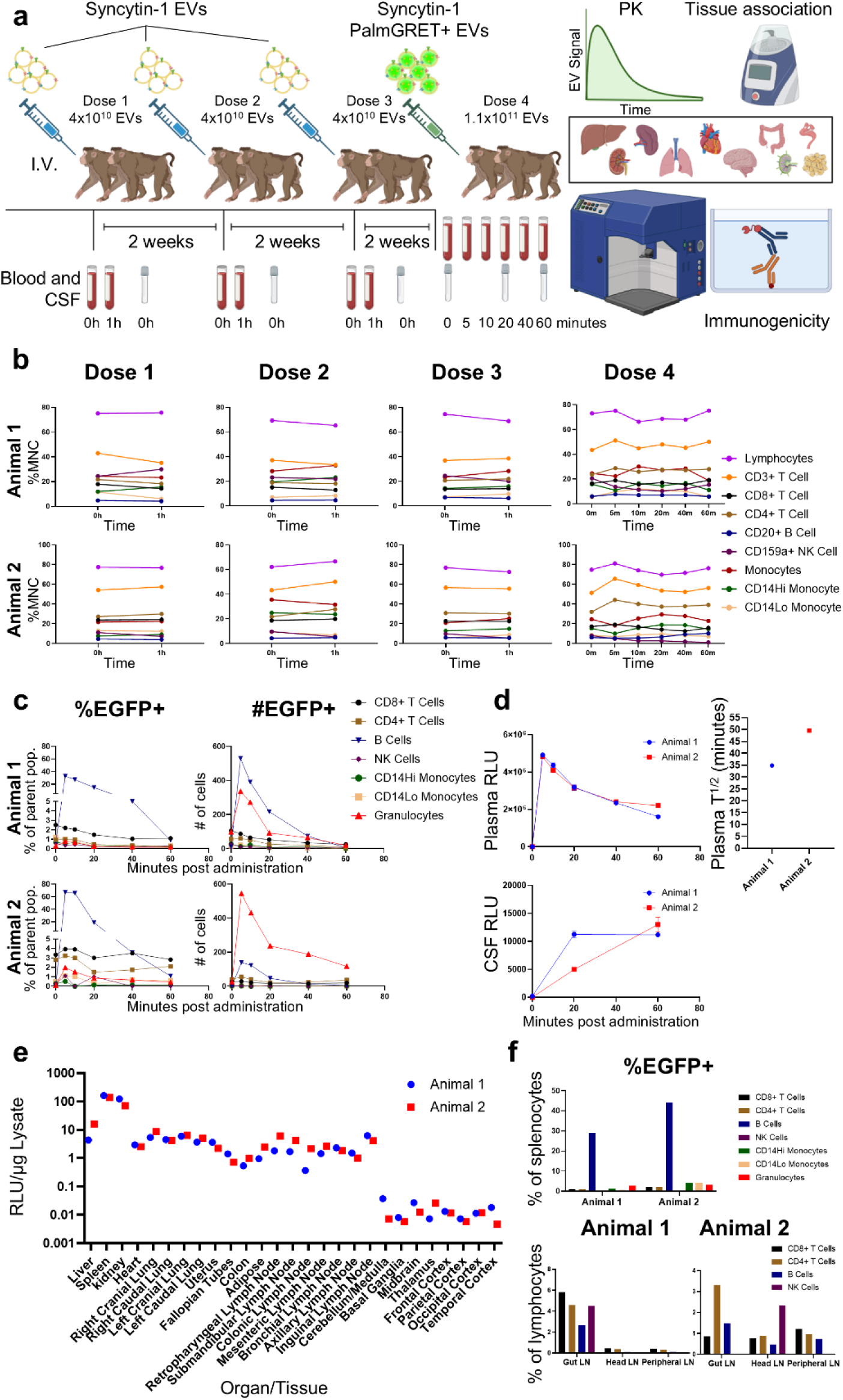
In vivo administration and tracking of Syncytin-1/PalmGRET EVs in non-human primates. (a) Schematic representation of the experimental workflow to introduce Syncytin-1 and Syncytin-1/PalmGRET EVs to pig-tailed macaques (M. nemestrina), intravenously. Four doses of EVs were administered with two-week periods between each dose. The final dose included a higher number of EVs and used Syncytin-1 EVs with the tracking molecule PalmGRET. Blood, cerebrospinal fluid (CSF), and organs/tissue were collected at designated time points and analyzed with a variety of methods to track EV biodistribution and monitor immunogenicity. (b) Line graphs representing the abundance of different blood cell subpopulations at different collection time points, across the four EV doses and two subject animals. Abundances are calculated as a percentage of the total mononuclear cell (MNC) population, as determined by flow cytometry gating. (c) Line graphs representing levels of Syncytin-1/PalmGRET EV interactions with blood cells, across different time points and two subject animals after EV dose 4. EV interactions were quantified as a percentage of the total cell type population (left panels) and raw magnitude of EGFP+ cells (right panels). (d) Line graphs (Left panels) representing detectable EV luminescence signal in blood plasma and CSF, over time after dose 4. Data points represent the mean relative light unit (RLU) signal of two technical replicates, while error bars represent standard deviation. The plasma data were used to calculate EV blood plasma half life (right panel). (e) Interleaved scatter plot representing EV accumulation and interaction with different organs and tissue lysates. Data points represent the mean RLU signal of two technical replicates, normalized to corresponding lysate protein concentration as determined by BCA. (f) Bar graphs representing EV association with immune cells derived from mechanically dissociated spleen (upper panel) and lymph nodes (lower panels). For all graphs, bars represent the EGFP positivity, determined as a percentage of the relevant total cell population, of different cell types.

Post-exposure, EV signal in blood declined over the course of one hour consistent with a one-phase exponential decay and a half-life of ∼35 and ∼50 minutes in the two subjects (average: ∼42 minutes)(Figure 5D, top panels). This was in contrast with our previous findings, in which unmodified EVs experienced accelerated blood clearance and half-lives of around ten minutes after four doses.^36^ Flow cytometry of cells in blood drawn at 5, 10, 20, 40, and 60 minutes after exposure suggested that Syncytin-1/PalmGRET EVs interacted strongly with B cells and granulocytes and less strongly with T cells and monocytes (Figure 5C). General PBMC interaction with Syncytin-1+ EVs was also confirmed by NanoLuc luciferase assays of bulk PBMCs (Supplemental Figure S4B).

EV signal also accumulated in cerebrospinal fluid (CSF) over the course of one hour post-IV administration (Figure 5D, bottom panel). Curiously, the CSF signal was several orders of magnitude higher than what we measured in a previous study^36^ of non-surface-modified PalmGRET EVs, even though the administered dose was lower in the current study. A hemoglobin ELISA was run on the CSF samples to check for blood contamination, which could lead to EV signal in CSF. The CSF with high EV signal also had increased blood contamination, offering a potential alternative explanation for the presence of EV signal (Supplemental Figure S4C).

### Tissue biodistribution: reduced liver and enhanced kidney accumulation

Luciferase signal was next measured in PBS-perfused solid tissues. For both subjects, signal per tissue mass was greatest in spleen and kidney, with concentration in both organs approximately ten-fold greater than in liver (Figure 5E). This finding is in stark contrast with previous EV biodistribution studies, including our own, which have reported high liver/spleen but low kidney accumulation.^21,31–36^ Levels similar to those in liver were observed in heart, lung, uterus, fallopian tubes, colon, adipose tissue, and various lymph nodes (with some apparent lymph node differences between the subjects). No signal was detected in any brain region for either subject.

Secondary immune organs (lymph nodes and spleen) were dissociated into single cell suspensions and assayed by flow cytometry. In the spleen, the majority of Syncytin-1+ EV interacting cells were B cells, matching blood findings, while monocytes, granulocytes, and T cells had very low levels of EV interaction (Figure 5F, top panel). Of gut, head/neck, and periphery lymph nodes, gut had the strongest evidence of EV-cell interaction as measured by flow cytometry, despite minimal signal in bulk tissue analysis (Figure 5F, bottom panels). In one subject, CD8+ T cell association predominated in gut lymph nodes, followed closely by CD4+ T cells and B cells; in the other, CD4+ T cells were followed by B cells and CD8+ T cells.

### Immunogenicity of intravenously administered Syncytin-1+ EVs

To understand EV immunogenicity, we examined innate and adaptive responses. Minimal or no changes were observed in plasma or CSF cytokine levels following EV administration (Figure 6A, Supplemental Figure S5). Any increases, e.g., for IL-6, were at or near the level of quantitation of the assay and/or within physiologic range. There were no consistent changes in total IgG, despite small fluctuations at different timepoints taken the same day (Figure 6B, right panel). EV-specific antibody ELISAs detected an increase in EV-specific IgG only in one animal and at dose 4, 6 weeks after the first Syncytin-1+ EV administration (Figure 6B, left panel). In contrast, in our previous study with unmodified EVs, EV-specific IgG was detected as early as the third dose.^36^ Additionally, the dilution used in that study was ten-fold greater than in the current work.

**Figure 6.**
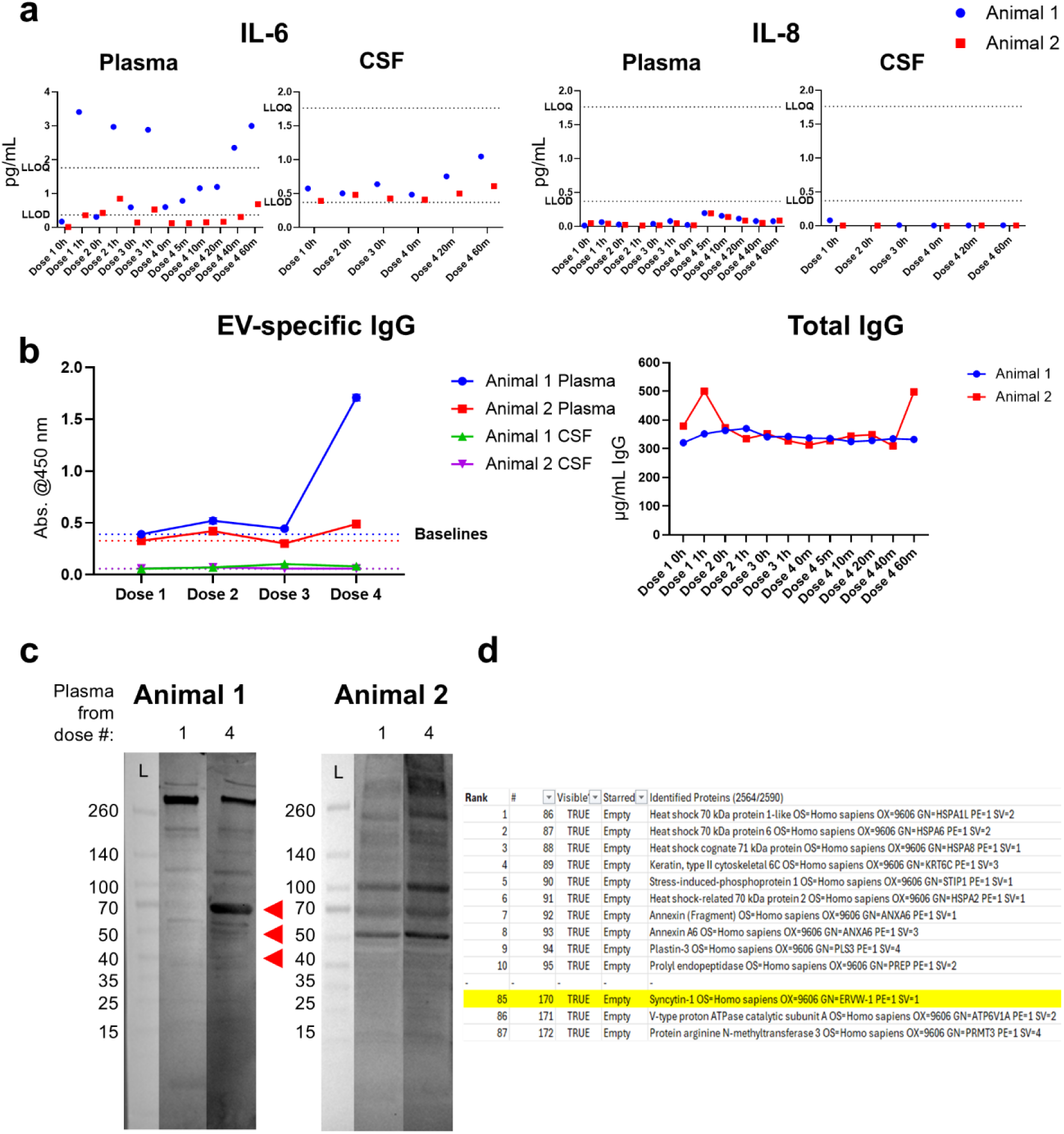
Immunogenicity of intravenously administered Syncytin-1 EVs. (a) Interleaved dot plots representing detection of inflammatory cytokines IL-6 and IL-8 in NHP plasma and CSF taken at various time points across the four EV doses for two subject animals. Data points represent interpolated mean cytokine concentrations of two technical replicates, as determined my multiplexed electrochemiluminescent ELISA. The hashed line labeled “LLOQ” represents the lower limit of quantitation for the cytokine; the hashed line labeled “LLOD” represents the lower limit of detection. (b) Line graphs representing the detection and quantification of EV-specific and total IgG in NHP plasma (both panels) and CSF (left panel only) taken at after each of the four EV doses for two subject animals, as determined by ELISA. (Left panel) Data points represent the mean absorbance at 450 nm wavelength of two technical replicates; error bars represent standard deviation. The colored hashed lines represent baseline absorbance values, determined by the values for dose 1 (naïve animals). (Right panel) Data points represent the mean interpolated total IgG concentration of two technical replicates. (c) Immunoblot analysis of Syncytin-1 EV lysates, using NHP plasma taken before dose 1 (naïve) and after dose 4 as a source of primary antibody for both subject animals. Red arrows indicate protein bands that appear after probing with EV-exposed plasma and not with naïve plasma, for animal 1. (d) Table displaying the most abundant proteins identified by gel-excision mass spectrometry of the major protein band appearing around 70 kDa (S70); equivalent immunoblot membrane location indicated by the top red arrow.

To identify which EV proteins were recognized, we probed electrophoresed Syncytin-1+ EV lysates with naïve (dose 1) and Syncytin-1+ EV-exposed plasma (dose 4, 6-weeks post 1^st^ EV administration). For the subject without an ELISA-positive response, the immunoblot banding pattern was identical, suggesting that any antibodies recognizing EV proteins preexisted in the subject (Figure 6C, animal 2). For the ELISA-positive subject, new immunoblot bands appeared for the exposed plasma: an intense band of ∼70 kDa (S70), with two lighter bands around 50 (S50) and 40 (S40) kDa (Figure 6C, animal 1). Gel excision mass spectrometry of the ∼70 kDa S70 band revealed common EV proteins like heat shock proteins, soluble cellular enzymes, and annexins, but Syncytin-1 was also detected, ranked as the 85^th^ most abundant protein when proteins from the blank gel or digest control were excluded (Figure 6D and Supplemental data 1). The 50 and 40 kDa bands also contained peptides derived from Syncytin-1 at low levels, representing the 120^th^ and 298^th^ most abundant protein, respectively (Supplemental data 1). These data suggest that Syncytin-1 may have provoked an immune response in one subject but also provide clues about native proteins that may elicit anti-EV adaptive immune responses.

## Discussion

This study reports a striking change in EV cellular interaction, biodistribution, and immunogenicity as a result of modification with the HERV envelope protein Syncytin-1. Three tested HERV envelope proteins^39–41^ were engineered onto the EV surface and did not significantly alter EV size. They were preferentially incorporated by EVs, consistent with established retroviral biology.^50^ HERV envelope proteins also tended to be in a mature and ready-to-fuse form^48^ in EVs versus cells. Retroviral envelope proteins are co-translationally inserted into the endoplasmic reticulum and transit through the trans-Golgi network (TGN), where they are cleaved by furin protease before incorporation into the plasma membrane.^51^ This suggests that HERV envelope+ EVs are ectosomes, although the presence of exosomes containing mature HERV proteins cannot be ruled out. The presence of both mature and immature HERV envelope proteins in soluble protein fractions from size exclusion chromatography could be due to increases in cell death with HERV envelope overexpression or to an unknown secretion pathway for HERV envelope proteins. However, EVs were successfully separated from this soluble fraction.

Our *in vitro*, *ex vivo*, and *in vivo* data suggest that the HERV envelope Syncytin-1 modulates EV tropism and cellular interaction. Modes of interaction include binding at the plasma membrane (without fusion) and internalization of EVs and/or cargo, the latter encompassing EV-membrane fusion and internalization of intact EVs. Syncytin-1+ EVs had 2-fold increased association but no increase in cellular uptake with embryonic kidney (HEK293T) cells, suggesting surface binding, while there was 6-fold higher association and 3-fold higher uptake with cervical adenocarcinoma (HeLa) cells. Different SLC1A5 (Syncytin-1 receptor) levels or metabolic activities are potential explanations. In NHP blood/PBMCs *ex vivo*, B cells had the highest levels of EV interactions regardless of surface modification (Syncytin-1, VSV-G, etc.), similar to our previous findings.^36,52,53^ However, EVs with viral surface proteins had globally reduced interactions with other PBMC subtypes, most prominently monocytes. Possibly, glycan-modified endogenous and exogenous viral proteins shield the EV from circulating opsonins or other factors, preventing clearance by the mononuclear phagocyte system (MPS).^54^

*In vivo*, Syncytin-1+ EVs displayed a large reduction in liver accumulation and a corresponding increase in kidney and CSF accumulation when administered intravenously to non-human primates (NHPs). Numerous rodent studies and a handful of NHP studies including our own have tracked EV biodistribution, consistently reporting EV clearance mostly in liver and spleen.^21,31–36^ We propose two complementary factors that may contribute to altered biodistribution. First, Syncytin-1 receptor SLC1A5 may drive local retention. For example, SLC1A5 is highly expressed in kidney proximal tubules^55^, and there is evidence that some EVs can cross the glomerular filtration barrier to reach glomerular podocytes^56^. SLC1A5 is also expressed in the brain near blood barriers^57^ and has been found in the rat choroid plexus.^58^ Second, glycan shielding^54^ and hence, reduced clearance by mononuclear cells and Kupffer cells of the liver may give EVs a better chance to find cognate receptors elsewhere.

The study yielded several interesting observations about EV-immune cell interactions in lymph nodes and spleen. Lymph node B cells did not associate with EVs to the same degree as their circulating and splenic counterparts, suggesting that organ architecture or surface characteristics of B cells in lymph nodes may influence EV interactions, or too few EVs reached lymph nodes compared to spleen. Alternatively, the high amounts of Syncytin-1+ EVs that accumulate in spleen may lead to higher chances of splenic B cell interaction in the marginal zone. Also, there was substantial intersubject variability in EV association with CD4+ and CD8+ lymph nodes in our study.

Syncytin-1+ EVs also had reduced immunogenicity and little evidence of accelerated blood clearance even after multiple doses. As demonstrated in our previous NHP studies^36^, repeated dosing of unmodified EVs leads to progressively reduced half-life in circulation, correlating with development of EV-specific antibody responses. EV-specific antibody responses were comparatively minimal or delayed for Syncytin-1+ EVs. Glycan shielding^54^ of viral envelope proteins on EVs might also prevent antibody development and recognition, further contributing to the longer half-life of Syncytin-1+ EVs (∼42 minutes).

Our study has several limitations. We compare results of *in vivo* administration of Syncytin-1+ EVs with a previous study of unmodified EVs.^36^ While the frequency of doses was the same, dosage amounts differed slightly between the studies and were lower, on average, here. However, our *in vitro* and *ex vivo* results support major differences between engineered and native EVs. Second, as for most EV reporters, we cannot rule out the possibility that PalmGRET has somehow escaped EVs. Even so, it is still important that PalmGRET (as a surrogate for therapeutic cargo) does show differences that correlate with EV modification. Lastly, although our study provides proof-of-principle for achieving altered behavior of EVs through engineering, the small number of subjects means that more work is needed, especially to understand outcomes that showed great variability.

Additional studies will be useful to answer several outstanding questions. With which kidney cells do Syncytin-1+ EVs interact? Is SLC1A5 the main determinant of this interaction? To what extent does glycan shielding influence immunogenicity, biodistribution, and reduced clearance of viral protein-displaying EVs? We did not test Syncytin-2 *in vivo*. Since the Syncytin-2 receptor MFSD2a is highly expressed on the blood brain barrier^59^, could Syncytin-2+ EVs be targeted to brain tissue, unlike Syncytin-1+ EVs? Importantly, can we replicate our results using non-invasive imaging techniques?^60^

In conclusion, we show that Syncytin-1+ EVs have altered tissue interaction, reduced PBMC association, increased half-life, and minimal immunogenicity. Several other HERV envelope proteins also show promise based on *in vitro* and *ex vivo* experiments. We hope that these findings will drive more research into harnessing endogenous retroviral elements for EV engineering.

## Methods

### Cell culture

Expi293F cells (Thermo Fisher Scientific) in Expi293 media (Gibco) in shaker flasks on an Orbi-Shaker CO_2_ (Benchmark Scientific) at 135 RPM (37°C, 8% CO_2_). HEK293T and HeLa cells (ATCC) were maintained in DMEM (Gibco, 4.5 g/L D-Glucose + L-Glutamine) with 10% heat-inactivated fetal bovine serum (FBS, R&D Systems) and 1% penicillin/streptomycin (Thermo Fisher Scientific). Cells were passaged using TrypLE (Thermo Fisher Scientific) at 80% confluence. All cells were used for only up to 30 passages.

### Plasmids

The pLenti-PalmGRET reporter plasmid^49^ was from C.P. Lai (Addgene #158221). Syncytin-1 and Syncytin-2 plasmids were a gift from J. Tilton. The HERV-K 108 Envelope plasmid was from Dr. L. Mulder. The VSV-G plasmid, pMD2.G, was from D. Trono (Addgene #12259).

### HERV envelope expression in HEK293Ts

HEK293Ts were plated into 6-well plates (250,000 cells in 2 mL) and transfected 24 hours later using JetOptimus reagent (PolyPlus). After 72 hours, cells were detached using TrypLE and lysed on ice with lysis buffer: 10 mL 1X PBS, 100 µL Triton-X 100, and 100 µL 100X HALT Protease and Phosphatase Inhibitor Cocktail (Thermo Fisher Scientific). After 15 minutes, whole cell lysates (WCLs) were clarified at 17,000g at 4°C. WCLs were stored at -20°C.

### EV production and separation

Expi293F cells were grown to >3×10^6^ cells/mL in volumes corresponding to flask size: 30 mL in a 125-mL flask, 60 mL (250), 250 mL (1L). Cells were transfected per manufacturer’s instructions using ExpiFectamine 293 (Thermo Fisher Scientific). ExpiFectamine Enhancers were added the next day to boost pDNA expression. 72 hours post-transfection, conditioned medium was cleared at 1000g for 20 minutes. Cell pellets were washed with PBS and lysed as previously described above. Supernatant was spun at 2000g for 20 minutes, and the 2K pellet was resuspended in 1 mL lysis buffer and stored at -20°C. EV-containing supernatant was pooled and concentrated to ∼20 mL by tangential flow filtration (TFF) using 100 kDa Vivaflow 50R cassettes (Sartorius) and a Cole-Parmer Masterflex L/S peristaltic pump. EVs were further concentrated to 10 mL using Amicon Ultra 15 mL 100 kDa centrifugal filters, spun at 4000g. For lower starting volumes EVs were concentrated to 0.5 mL by spin filters only). EV concentrate was loaded onto SEC columns (qEV10 70 nm or qEV Original Gen2 70 nm, Izon) and eluted with 1X PBS. 5 mL (qEV10) or 0.5 mL (qEV Original) fractions were collected. EV enriched fractions 1-4 were pooled and reconcentrated by centrifugal filter, then aliquoted and stored at - 80°C. EV purification and characterization were consistent with MISEV.^3^

### Immunoblotting

WCL protein concentration was determined by BCA assay (Pierce). 20 µg of WCL, 24 µL of 2K pellet and pooled EVs, and 24 µL of SEC fractions were combined with 5X Pierce Lane Marker Reducing Sample Buffer (Thermo Fisher Scientific) and heated for 10 minutes at 95°C. Samples were loaded into a 4-15% Criterion TGX Stain-Free precast gel (Bio-Rad) and electrophoresed (Polyacrylamide Gel Electrophoreses: PAGE) at 100V. After total protein imaging (Gel Doc EZ Imager, Bio-Rad), proteins were transferred to PVDF membranes using an iBlot2 (Thermo Fisher Scientific). Membranes were blocked using 5% Blotting-Grade Blocker (Bio-Rad) in PBS + 0.1% Tween-20 (PBST) for 1 hour at RT and incubated overnight at 4°C with primary antibodies: rabbit anti-Syncytin-1 (1:500, ab139693, Abcam), rabbit anti-Syncytin-2 (1:1000, ab230235, Abcam), mouse anti-HERV-K 108 Envelope (1:1000, HERM-1811-5, AMSbio), rabbit anti-TSG101 (1:500, ab125011, Abcam), rabbit anti-Syntenin-1 (1:1000, ab133267, Abcam), rabbit anti-Calnexin (1:1000, ab22595, Abcam), mouse anti-HA Tag (1:1000, #65738, Cell Signaling), and mouse anti-EGFP (1:1000, ab184601, Abcam). Blots were washed and incubated for 1h at RT with anti-mouse IgGκ-BP-HRP (1:5000, sc516102, Santa-Cruz Biotechnology) or anti-rabbit IgG-HRP (1:5000, sc2357, Santa-Cruz Biotechnology) secondary antibodies. Blots were washed again and developed with SuperSignal West Pico PLUS or Femto Maximum Sensitivity substrate (Pierce). Chemiluminescence was detected by iBright FL1000 (Thermo Fisher Scientific).

### Nanoflow cytometry

EVs were diluted 1:10 to 1:1000 in 1X PBS for measurement by Flow Nanoanalyzer (NanoFCM), pre-calibrated for concentration with 250 nm silica nanoparticles (NanoFCM, #QS2503) and sizing with a silica nanoparticle cocktail (NanoFCM, #516M-Exo). Particle concentration, size distribution, and EGFP positivity (%) were measured.

### Transmission electron microscopy (TEM) and immunogold TEM

EVs were diluted to 1×10^10^ particles/mL in 1X PBS. 10 µL was adsorbed to glow-discharged carbon-coated 400 mesh copper grids (EMS) by flotation for 2 minutes. Grids were blotted, rinsed with Tris-buffered saline (TBS), and negatively stained with 1% uranyl acetate in deionized water with tylose. After blotting and aspiration to form a thin stain layer, grids were imaged on a Hitachi H-7600 TEM operating at 80 kV with an AMT XR80 CCD camera.

For immunogold, EVs were fixed in 2% paraformaldehyde, adsorbed to copper grids (EMS) for 2 minutes, rinsed in filtered 1X TBS (Tris-buffered saline) before incubating in 0.05 M glycine, then blocked in filtered 5% milk/TBS for 30 min. Antibodies [rabbit anti-Syncytin-1 (1:500, ab139693, Abcam), rabbit anti-Syncytin-2 (1:1000, ab230235, Abcam), mouse anti-HERV-K 108 Envelope (1:1000, HERM-1811-5, AMSbio] were diluted 1:30 in TBS/0.01% BSA and incubated on grids in a humidity chamber (2 hours, RT). After washing in TBS for 50 min, grids were incubated with 6 nm gold-conjugated anti-mouse or anti-rabbit antibodies (Jackson Immuno, 1:10 in TBS). Grids were washed in TBS for 10 min, fixed in 2% glutaraldehyde for 5 min, rinsed in TBS, negative stained on 2 consecutive drops of filtered aqueous 1% Uranyl Acetate for 30 seconds each (Pella), blot dried with Whatman # 1 filter paper, and imaged as above.

### Molecular cloning

Primers were designed with Benchling to add a 2x HA tag sequence to the C-terminus of each HERV envelope (Syncytin-1, Syncytin-2). Gibson assembly was performed with 2X NEBuilder HiFi DNA Assembly Master Mix (New England Biolabs) and 100ng of each PCR product. Products were electroporated into ElectroMAX Stbl4 Electrocompetent *E. coli* (Thermo Fisher Scientific) using a MicroPulser (Bio-Rad), and bacteria were grown on Ampicillin LB plates. pDNA was isolated from bacterial clones and sequence verified.

### *In vitro* EV association and uptake assays (NanoLuc luciferase)

HEK293Ts (50,000 cells) or HeLa cells (20,000 cells) were plated into 96-well flat-bottom plates in 100 µL DMEM. Next day, 70 µL was removed from each well, and 1000 PalmGRET EVs/cell were added in 100 µL fresh DMEM. After 6 hours, EVs/media were removed, and cells were washed 3x with 100 µL PBS. For EV association assays, cells were lysed for 5 minutes with 100 µL NanoGlo Luciferase assay reagent (Promega) and moved to white 96-well plates (Corning). For EV uptake assays, cells were detached with 50 µL of 1X 0.05% Trypsin-EDTA (Gibco) in calcium/magnesium-free PBS (5 min, 37°C). After trypsin neutralization (150 µL DMEM), cells were moved to 96-well V-bottom plates (Corning), washed 2x in PBS, resuspended in 100 µL assay reagent, and moved to white 96-well plates. As a control, EV NanoLuc luciferase activity was measured by adding the same quantity of EVs directly to the white plate. Luminescence was measured on a SpectraMax ID5 (Molecular Devices). %Association and %Uptake values were calculated by dividing the cellular relative light unit (RLU) signal by the average control EV-only RLU signal (3 wells) and multiplying by 100. Data were normalized to unmodified PalmGRET EVs.

### Confocal microscopy

A Nunc MicroWell 96-Well Flat-Bottom Optical Cover-glass (1.5) Microplate (Thermo Fisher Scientific) was coated with 100 µL 4 µg/mL poly-L-ornithine (Millipore Sigma) in cell culture grade water (Quality Biological; 3 hours, 37°C), rinsed with PBS, and coated (overnight, 37°C) with 100 µL 4 µg/mL laminin (Millipore Sigma) in HBSS (Gibco). After laminin removal and PBS rinse, 15,000 HEK293Ts were plated in 100 µL DMEM. The following day, cells were treated with 1000 EVs/cell as above. After 6 hours, cells were washed 3x with 100 µL PBS, fixed with 4% paraformaldehyde (EMS) in PBS (15 min), and washed with 100 µL 50 mM glycine (Millipore Sigma) in PBS (10 min). Cells were permeabilized with 100 µL 0.2% Triton-X 100 (Millipore Sigma) in PBS (10 min), blocked with 100 µL 1% BSA (w/v) in PBS (1 hour), and incubated (o/n, 4°C) with 100 µL 1 µg/mL rabbit anti-EEA1 (Cell Signaling Technology) and 1X phalloidin-647 (Abcam) in 1% BSA. Cells were washed and incubated with 100 µL 1 µg/mL donkey anti-rabbit IgG PE (Biolegend) in 1% BSA. After 1 hour, nuclei were stained with Hoechst 33342 Ready Flow (Thermo Fisher Scientific) in PBS for 10 min, 100 µL PBS was added, and the plate was imaged using a Zeiss 880 Airyscan FAST confocal microscope: 63x magnification, Zen 2.3 SP1 software (Carl Zeiss AG). Channels for DAPI (410-450 nm; gain master=850), A488 (PalmGRET/EGFP)(490-544 nm; gain master=850), A568 (EEA1/PE)(561-583 nm; gain master=850), and A647(Phalloidin) (633-672 nm; gain master=850) were used with 5.0% laser intensity. Z-stacks were acquired with a 40.3 (1 AU) pinhole and 0.2 µm intervals, using digital refractive index correction=1. Images were averaged twice and obtained with speed=8 and 16-bit depth. Acquisition conditions were maintained across all groups. Image handling and processing for orthogonal views used FIJI (ImageJ). Colocalization analysis used the FIJI plugin JACoP.^61^ Pearson’s correlation values were determined using the same threshold values for all images: C2 (EEA1/PE)=4369 and C3 (PalmGRET/EGFP)=1460.

### *Ex vivo* EV blood spike-in assay

Blood was collected into tubes containing 100 µL ACD. 1 mL of anticoagulated blood in 2 mL Protein LoBind Eppendorf tubes was mixed with 1×10^9^ PalmGRET EVs (or PBS) on a Roto-Therm Mini Plus incubator (Benchmark Scientific) for 30 minutes at 37°C. Blood was diluted to 2 mL in 1X HBSS/EDTA, layered onto 4 mL Percoll (Cytiva) solution in SepMate-15 tubes (Stemcell Technologies), and centrifuged at 1200g for 10 minutes at RT to obtain a PBMC interphase. After washing and RBC lysis with ACK lysis buffer (Quality Biological), PBMC pellets were resuspended in 200 µL 1X PBS + 2% FBS and evaluated for EGFP signal with flow cytometry. %EGFP positivity data of cells were first normalized to account for differences in the %EGFP positivity of different EV conditions as follows:

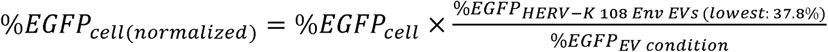

Averaged data from three independent experiments were normalized to unmodified EVs to calculate fold change.

### *In vivo* administration to macaques

Adult pigtail macaques (*M. nemestrina*, female, 11 and 13 years old) from the JHU pigtail colony received saphenous vein injections of 4×10^10^ Syncytin-1+ EVs in 1 mL PBS under ketamine sedation (10 mg/kg body weight) every two weeks for three exposures. For a fourth and final exposure, 1.1×10^11^ EVs were administered in 3 mL PBS. At indicated timepoints, blood was collected into ACD tubes and analyzed by flow cytometry and for complete blood counts. Plasma and PBMCs were also separated using Percoll in SepMate-50 tubes (Stemcell Technologies; 1200g, 10 min, RT). Plasma was stored at -80°C. After RBC lysis and 2x PBS washes, PBMCs were resuspended in HBSS, lysed, and stored at -20°C. CSF was collected via the cisterna magna and centrifuged at 2000g before storage at -80°C. After the final dose and biofluid collections, macaques were deeply anesthetized with ketamine and euthanized using IV pentobarbital before perfusion with PBS. Organs were removed and snap-frozen at -80°C. Portions of spleen and lymph nodes were processed directly for flow cytometry (see below).

### Flow cytometry and complete blood counts (CBCs)

100 µL whole blood or 200 µL isolated PBMCs were added to a cocktail of antibodies for 20 min at RT: mouse anti-CD159a-PE (Beckman Coulter), mouse anti-CD4-PerCP/Cy5.5 (BD Biosciences), mouse anti-CD20-e450 (Thermo Fisher Scientific), mouse anti-CD3-V500 (BD Biosciences), mouse anti-CD8a-BV570 (Biolegend), and mouse anti-CD14-BV650 (BD Biosciences). RBCs were lysed, and PBMCs washed 1X with PBS and resuspended in 500 µL PBS. Cells were measured with a BD LSR Fortessa. Cells were isolated from fresh spleen and lymph nodes using 18-gauge needle shearing in cold RPMI and a 100 µm cell strainer (Corning). RBCs in spleen tissue were lysed using RBC lysis buffer as before. 1×10^6^ splenocytes or lymphocytes were resuspended in 100 µL PBS/2% FBS and stained as for PBMCs. CBCs were obtained from whole blood using a Procyte Dx (IDEXX).

### Organ and Tissue Homogenates

50 mL N-PER neuronal protein extraction reagent (for brain) or T-PER tissue protein extraction reagent (Thermo Fisher Scientific) were mixed with 1 cOmplete Protease Inhibitor Cocktail tablet (Roche) for tissue lysis. ∼100 mg snap-frozen organs and tissues were cut and transferred to FastPrep Lysing Matrix D tubes (MP Biomedicals) on ice containing 1 mL of N-PER or T-PER. Homogenates were prepared using a FastPrep-24 homogenizer (MP Biomedicals) and centrifuged at 10,000g for 5 min to pellet insoluble debris. 700 µL supernatant (homogenate) was collected and stored at -80°C.

### Bichinchoninic assay (BCA)

PBMC lysates and organ/tissue homogenates diluted 1:10 in PBS were added to 200 µL BCA working reagent (Pierce) in 96-well plates. BCA absorbance was measured at 570 nm using an iMark (Bio-Rad). Protein concentrations were interpolated from a blank-subtracted standard curve.

### NanoLuc luciferase assay

White 96-well plates were loaded with 50 µL biofluids or lysates/homogenates per well, mixed with 50 µL NanoGlo Luciferase assay reagent (Promega). Bioluminescence was measured by SpectraMax ID5 (Molecular Devices). RLU values were normalized to protein concentration where applicable.

### Hemoglobin ELISA

CSF was thawed and serially diluted (1:20, 1:200, 1:2000) using 1X diluent (Monkey Hemoglobin ELISA, Abcam) following manufacturer’s protocol. Absorbance was measured at 450 nm by iMark Microplate Absorbance Reader (Bio-Rad) and interpolated onto a standard curve.

### Cytokine analysis

60 µL thawed biofluids were mixed with 60 µL diluent 2 (V-PLEX Proinflammatory Panel 1 NHP Kit, Mesoscale), and 50 µL was then added in duplicate to an ELISA plate following manufacturer’s instructions. Readings were done with Mesoscale Diagnostics MESO QuickPlex SQ 120MM. Standard curves were fitted using a 4-PL sigmoidal curve with 1/Y^2^ weighing.

### EV-specific antibody/antigen analysis

For total plasma IgG measurement, a 1:70,000 dilution was used with the Human IgG ELISA kit (Abcam)(cross reactive with NHP) following manufacturer’s instructions and per the Hemoglobin ELISA protocol above.

A modified version of our EV-specific ELISA^62^ was prepared using 2×10^10^ Syncytin-1+ EVs in 10 mL PBS. After coating the plate with EVs overnight, biofluid samples were diluted 1:100 with 1X Assay Buffer (Thermo Fisher Scientific), and 100 µL was added to each well in duplicate. 450 nm absorbance was read as above.

To measure binding of antibodies in plasma to EV proteins, 24 µL of Syncytin-1+ EVs (2.39×10^11^ EVs/mL) were mixed with 6 µL of 5X Pierce Lane Marker Reducing Sample Buffer (Thermo Fisher Scientific) and subjected to PAGE. For blotting, proteins were transferred to PVDF. After blocking, membranes were sequentially probed (1 hour each) with plasma diluted 1:1000 in 0.1% PBST, followed by goat anti-monkey IgG H&L HRP (Abcam) secondary antibody (1:5000), and imaged as previously. Alternatively, for proteomics, the gel was washed with DI H_2_O, stained for 1 hour with SimplyBlue SafeStain (Thermo Fisher Scientific), washed overnight, and imaged on iBright FL1000 (Thermo Fisher Scientific). The gel was then transferred to an acetonitrile-rinsed petri dish, and protein bands were excised. Gel slices were washed 2X with 50% HPLC-grade ethanol in water (Fisher) and dried.

### In-gel digestion

Gel pieces were twice destained in 50% methanol containing 50 mM dithiothreitol (DTT) at 60°C for 10 min. Proteins were reduced with 50 mM DTT at 60 °C for 1 h. Cysteine residues were alkylated with 100 mM iodoacetamide for 15 min at RT in the dark. Gel pieces were washed twice with 10 mM triethylammonium bicarbonate (TEAB) and dehydrated with 100% acetonitrile (ACN) before additional dehydration with 100% ACN. Gel pieces were rehydrated on ice with 25 µL trypsin (0.01 µg/µL) in TEAB solution for 15 min. An additional 75 µL of 10 mM triethylammonium bicarbonate was added. Digestion was performed overnight at 37 °C with shaking. Peptides were extracted sequentially for 20 min at RT with 60% ACN containing 0.1% trifluoroacetic acid twice, followed by extraction with 100% ACN. Supernatants were combined and dried using a Savant SPD121P SpeedVac concentrator (Thermo Fisher Scientific). Dried peptides were reconstituted in 0.1 formic acid, desalted using Oasis cartridges, and dried again.

### Mass Spectrometry

Peptides were analyzed by reverse-phase chromatography coupled to tandem mass spectrometry using an EasyLC nano HPLC system interfaced with an Orbitrap Fusion Lumos mass spectrometer (Thermo Fisher Scientific). Peptides were separated on an in-house packed 75 µm ID × 15 cm analytical column packed with ReproSIL-Pur-120-C18-AQ resin (2.4 µm, 120 Å, Dr. Maisch) using a 100 min linear gradient from 2% to 90% solvent B (0.1% formic acid in acetonitrile; solvent A: 0.1% formic acid in water) at a flow rate of 300 nL/min. MS1 spectra were acquired in the Orbitrap in positive ion mode over an m/z range of 375–1500 at a resolution of 120,000. Data-dependent acquisition was performed with a cycle time of 3 s using quadrupole isolation (1.2 m/z window) with a dynamic exclusion of 30 s. Precursors with charge states of 2–6 were selected for fragmentation by higher-energy collisional dissociation (HCD; normalized collision energy 32), and MS2 spectra were acquired in the Orbitrap at a resolution of 30,000.

The MS/MS data obtained from LC-MS/MS analyses underwent a search against the UniProt human protein database, comprising Swiss-Prot and TrEMBL entries released in 2024 and common contaminant proteins. The search was conducted using the Mascot algorithm within the Thermo Proteome Discoverer software package (version 3.1.1.93, Thermo Fisher Scientific). The search parameters included trypsin as the designated protease, allowing for a maximum of one missed cleavage. Precursor mass (MS1) and fragment mass (MS2) tolerances were both set at 10 ppm. Deamidation (NQ) and oxidation (M) were included as dynamic modifications, and carbamidomethylation of cysteine (C) was applied as a static modification. False discovery rate (FDR) filtering was applied at 1% for both peptides and proteins using the Percolator node and Protein FDR Validator node, respectively.

### Statistics and EV half-life calculation

Statistical differences were assessed by one-way ANOVA with Dunnett’s correction for multiple comparisons using GraphPad Prism 10.4; differences with p<0.05 were considered statistically significant. EV half-life in plasma was determined by log-transforming NanoLuc luciferase values and plotting them versus time on the X-axis in a log-lin chart. Linear regression was performed to obtain a slope, which was used to calculate half-life using the equation: T_1/2_ = log(2)/slope.

### IACUC Statement

All procedures involving animals were reviewed and approved by the Institutional Animal Care and Use Committee of Johns Hopkins University (IACUC protocol number: PR24M193).

### Data sharing, availability, and reporting frameworks

The data used for this study are available within the article, in supplementary files, or available upon request to the corresponding author. Experimental details have been submitted to the EV-TRACK database and are available under ID: EV260028. Flow cytometry experiments involving EV detection were performed following the MIFlowCyt-EV guidelines and reporting framework.

## Author Contributions (CRediT)

Conceptualization: ZT, KWW

Methodology: ZT, MS, ES, OG, MF, BP, NC, IB, SQ, KWW

Validation: ZT Formal analysis: ZT

Investigation: ZT, MS, ES, OG, SM, MF, BP, NC, IB, SQ

Resources: ZT, ES, SQ, JM, KWW

Writing – original draft: ZT

Writing – review and editing: ZT, KWW

Visualization: ZT

Funding acquisition: ZT, KWW

## Acknowledgements

The authors would like to thank Barbara Smith and the Johns Hopkins University School of Medicine microscope core facility for assistance with electron microscopy and confocal microscopy experiments. We would additionally like to thank Dr. Chan-Hyun Na and the Center for Proteomics Discovery core for assistance with mass spectrometry. We also thank Dr. Charles Lai for the PalmGRET plasmid, Dr. John Tilton for the Syncytin-1 and Syncytin-2 plasmids, Dr. Lubbertus Mulder for the HERV-K 108 Envelope plasmid, and Dr. Didier Trono for the VSV-G plasmid. Importantly, we would like to thank all members of the Witwer Laboratory, the Retrovirus Laboratory, and the Johns Hopkins Research Animal Resources for assistance with various aspects of the research project. Lastly, we would like to thank Dr. Joy Wolfram and Dr. Mona Batish for their guidance and collaboration on the sponsored project.

---

**Figure S1.**
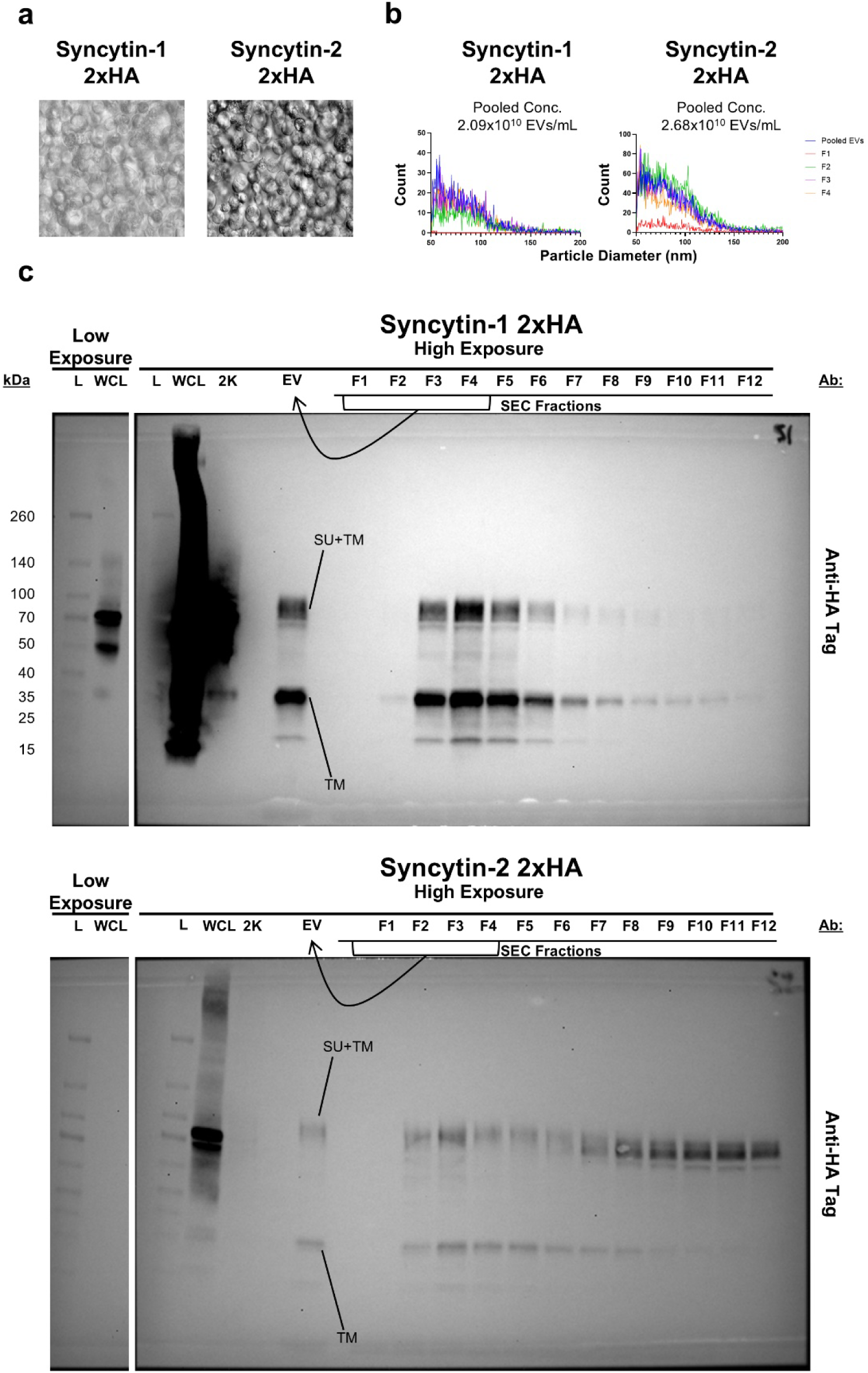
Cell and EV distribution of mature and immature HERV envelope proteins. (a) Brightfield microscope images of Expi293F cells transfected with HERV envelopes engineered to carry 2xHA tags at their C-terminus. (b) Nanoflow cytometry plots showing the size distribution of 2xHA tagged HERV envelope EVs derived from Expi293F cells. (c) Immunoblot analysis of protein ladder (L), whole cell lysate (WCL), 2000g (2K) pellet, pooled EVs, and size exclusion chromatography (SEC) fractions from Expi293F cells transfected with 2xHA tagged HERV envelopes. Bands corresponding to the full-length protein or furin cleavage products are indicated by labels. A low exposure panel is used to show the distribution of Syncytin-1 protein and cleavage products in the WCL.

**Figure S2.**
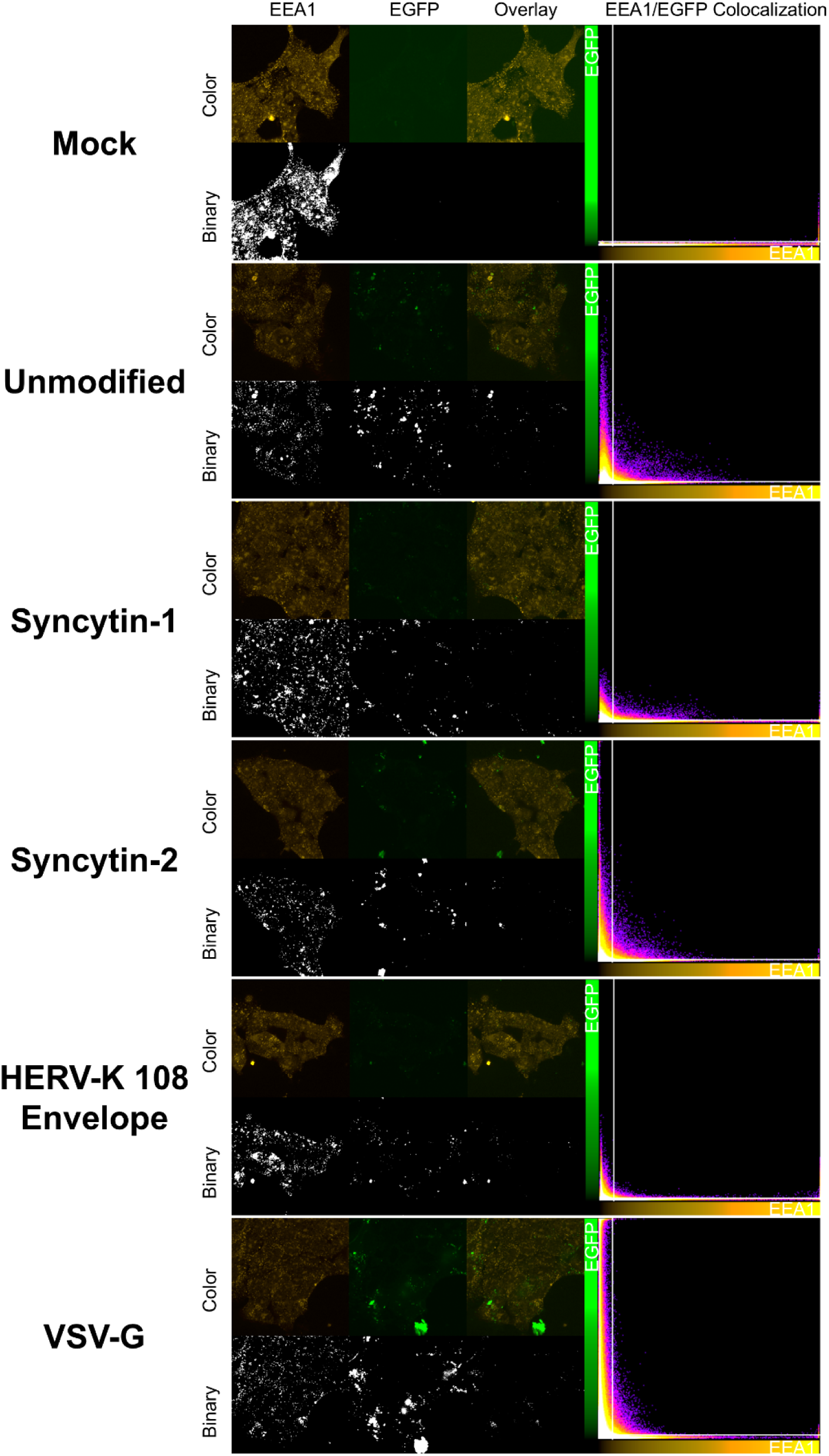
Confocal analysis of PalmGRET/HERV envelope EV uptake and endosomal escape in HEK293T cells. Confocal microscopy images (63X magnification) of HEK293T cells after incubation with different types of PalmGRET EVs. Cells were stained with antibodies or fluorescent moieties to identify endosomes (EEA1-PE), nuclei (Hoechst 33342 – not visible), and cytoskeletal actin (Phalloidin-647 – not visible). Colocalization of green (EGFP+ EVs) and orange (EEA1+ endosomes) pixels was assessed to crudely determine EV colocalization or escape from endosomes and quantified using Pearson correlation. Images were adjusted by increasing brightness by 40% and lowering contrast by 40% to better visualize fluorescence.

**Figure S3.**
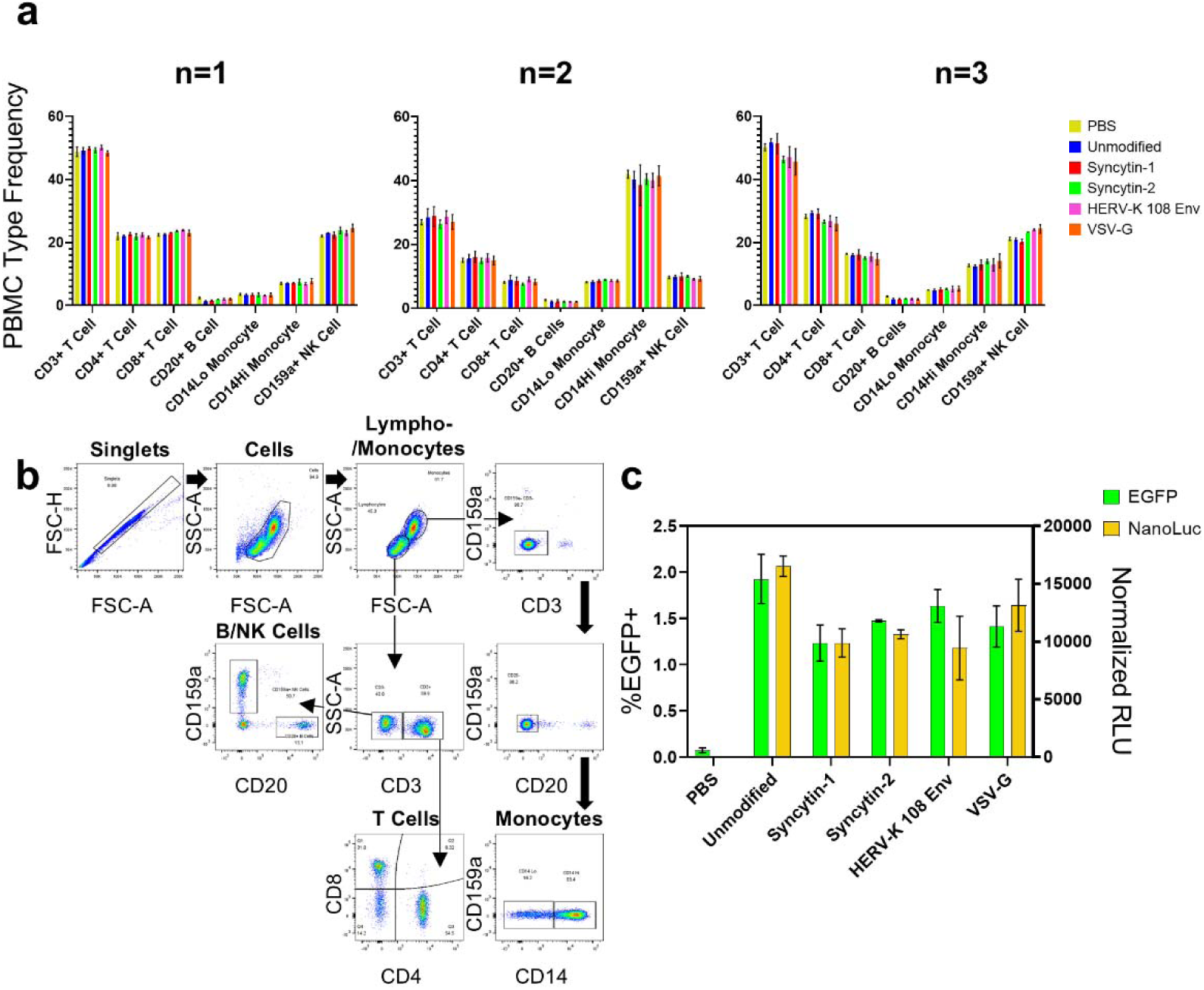
Assay to measure EV interaction with NHP blood cells *ex vivo*. (a) Bar graphs representing the prevalence of various peripheral blood mononuclear cell (PBMC) subtypes, as measured by percent of total PBMCs (top panels) and numerical quantity (bottom panels), after mixing with different PalmGRET/HERV envelope EV conditions and across independent biological experiments. Bars represent the mean of 3 technical replicates; error bars represent the standard deviation. (b) Flow cytometry staining and gating scheme used to identify and separately analyze different PBMC subpopulations. (c) Double Y-axis bar graph comparing the results of two techniques designed to measure PalmGRET EV interactions with NHP PBMCs, across different EV conditions. Bars represent mean %EGFP positivity (green) or mean bulk PBMC lysate luminescence (yellow), measured using relative light units (RLUs). RLU values were normalized to the protein concentration of the lysate, as determined by bicinchoninic acid (BCA) assay.

**Figure S4.**
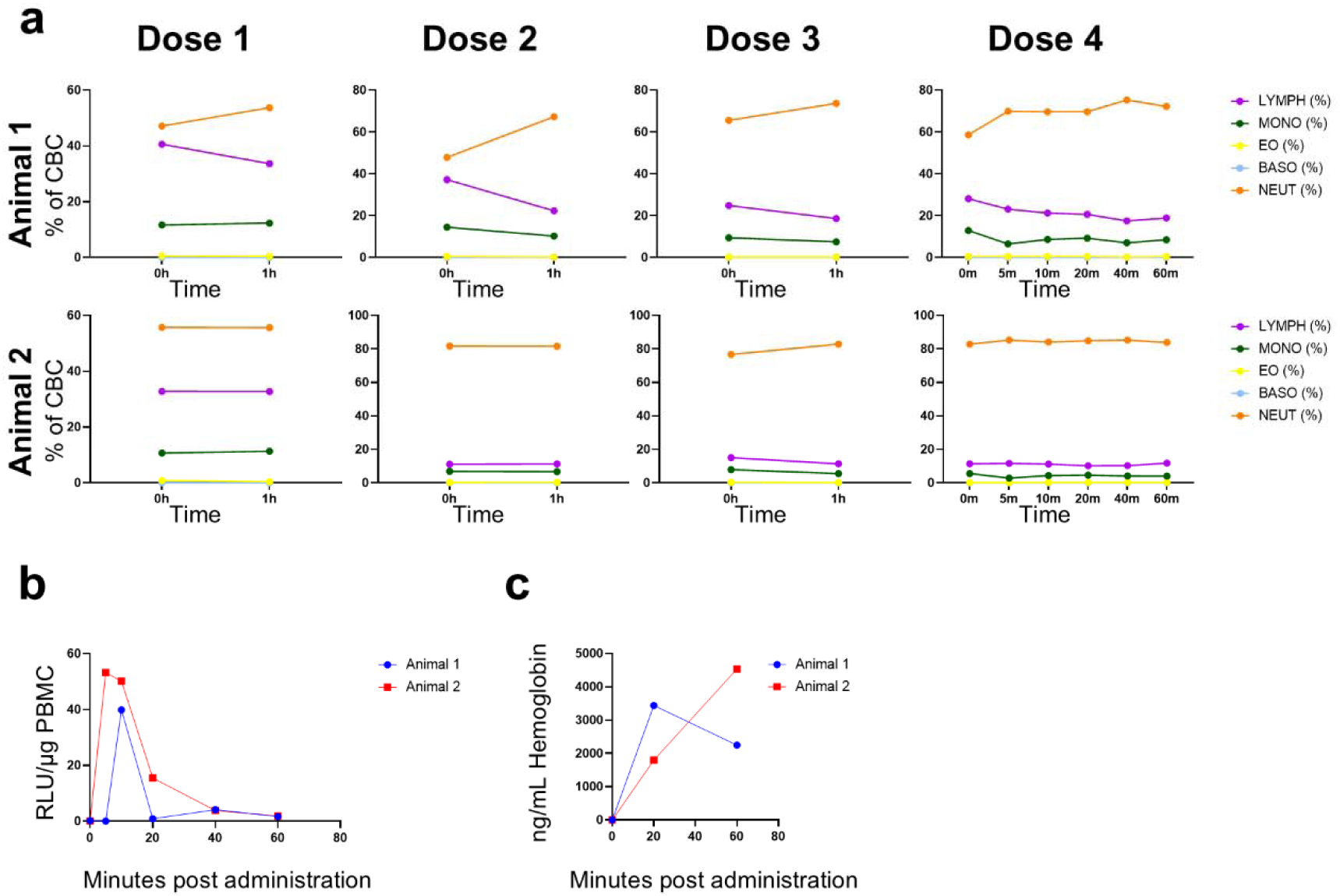
Additional analysis of blood and CSF from *in vivo* experiments. (a) Complete blood counts (CBCs) of whole blood taken at various time points, across the four EV doses and two subject animals. Data are calculated as a percentage of the total blood cell population, as determined by a combination of light scattering and electrical impedance. (b) Line graph representing EV interaction with bulk PBMCs taken at different time points after EV dose 4, as determined by luminescence via luciferase assay. Data points represent the mean RLU values normalized to the PBMC lysate protein concentration as determined by BCA assay. (c) Line graph representing presence of hemoglobin in CSF taken at different time points after EV dose 4, as determined by NHP hemoglobin ELISA. Data points are the mean interpolated hemoglobin concentration of two or three serial dilutions of the same CSF sample, each assayed in technical duplicate.

**Figure S5.**
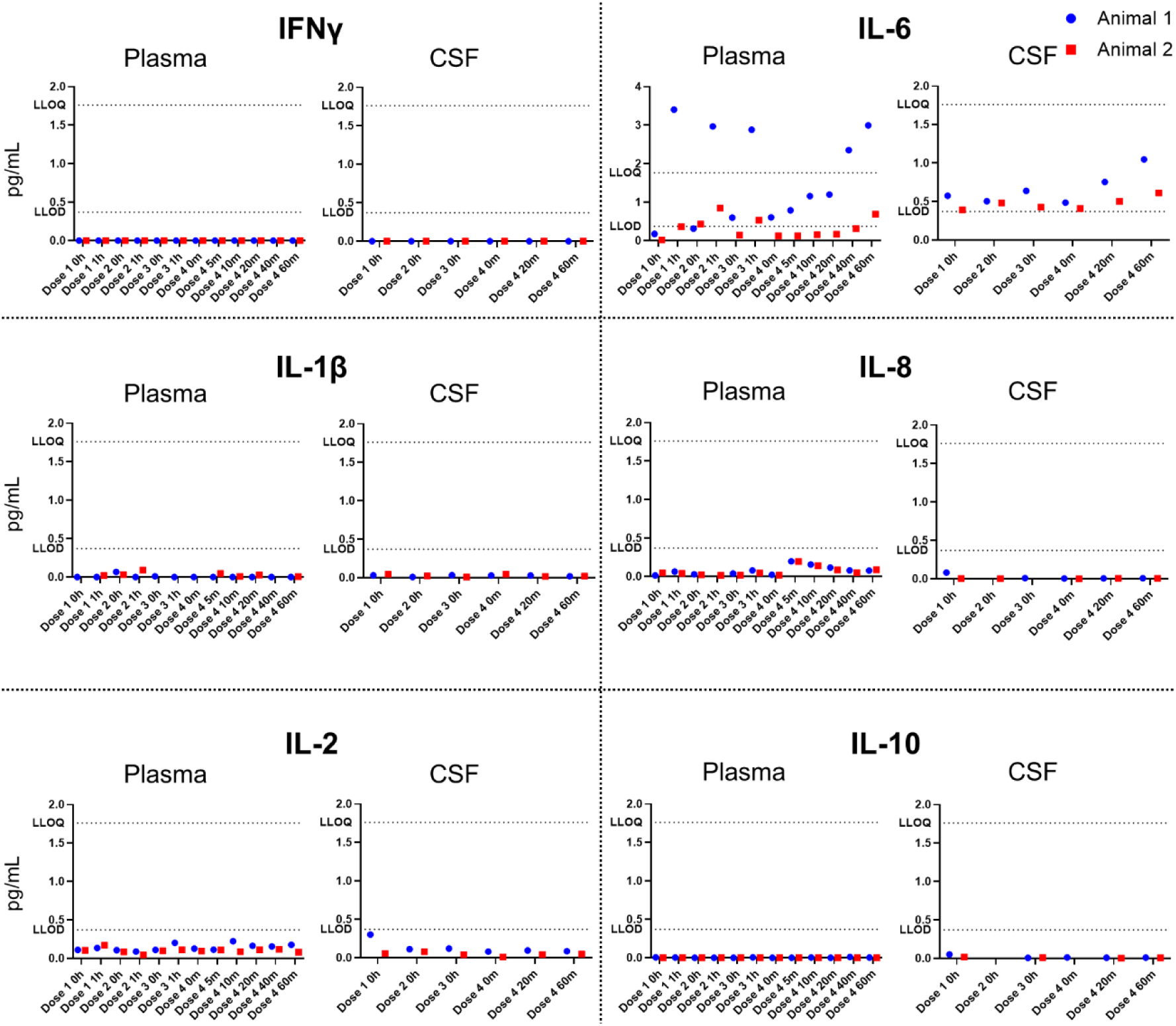
Innate immunogenicity of intravenously administered Syncytin-1 EVs. Interleaved dot plots representing detection of various inflammatory cytokines in NHP plasma and CSF taken at various time points across the four EV doses for two subject animals. Data points represent interpolated mean cytokine concentrations of two technical replicates, as determined by multiplexed electrochemiluminescent ELISA. The hashed line labeled “LLOQ” represents the lower limit of quantitation for the cytokine; the hashed line labeled “LLOD” represents the lower limit of detection.

